# Triceps surae and Achilles tendon contributions to ankle stiffness depend on movement state

**DOI:** 10.64898/2026.08.19.745788

**Authors:** Kristen L. Jakubowski, Daniel Ludvig, Eric J. Perreault, Sabrina S.M. Lee

## Abstract

Ankle stiffness is decreased during movement compared to posture; however, the etiology of this decrease remains unknown. Determining what gives rise to this decrease is critical for understanding how humans successfully interact with their physical world and how that ability is compromised by functional impairments. While the triceps surae and Achilles tendon primarily dictate ankle stiffness, the relative contributions across posture and movement remain unknown. Therefore, our study sought to quantify the relative contributions of the muscle and tendon to ankle stiffness and how those contributions differ between posture and movement. We used our technique, which combines B-mode ultrasound imaging with joint-level perturbations, to quantify ankle, muscle, and tendon stiffness simultaneously. Since ankle, muscle, and tendon stiffness all scale with torque, participants matched torque between posture and movement tasks. During posture, the Achilles tendon is the dominant contributor to ankle stiffness. However, during movement, the triceps surae and Achilles tendon contribute more equally to ankle stiffness, which can be attributed to a significant decrease in muscle stiffness during movement. Here, we provide the first empirical data on how state-dependent properties of the triceps surae and Achilles tendon contribute to ankle stiffness in conditions relevant to locomotion.

**SUMMARY STATEMENT:** Decreased triceps surae stiffness during movement leads to shared contributions from the Achilles tendon and triceps surae to ankle stiffness, unlike postural stiffness driven primarily by the Achilles tendon.

## INTRODUCTION

Humans perform a wide range of locomotor tasks safely and economically: seamlessly transitioning from posture to movement, navigating diverse terrains, and responding to unexpected postural disturbances. This is achieved partly by appropriately regulating ankle stiffness—the dynamic relationship between an imposed displacement and the torque generated in response [1–3]. Under active conditions, the primary contributors to ankle stiffness in the sagittal plane are the triceps surae muscles and Achilles tendon. While we recently quantified the contributions from the triceps surae and Achilles tendon to ankle stiffness during conditions relevant to stance [4], it remains unclear how the triceps surae and Achilles tendon contribute to ankle stiffness during movement tasks, such as locomotion. It is well documented that stiffness decreases during movement compared with posture at the ankle [5] and other joints [6, 7]. Although there are clear differences in muscle and tendon kinematics between posture and movement, the etiology of the decreased ankle stiffness during movement has not been identified. A clear understanding of the respective roles of the triceps surae and Achilles tendon between posture and movement will advance our fundamental understanding of how humans meet the wide range of locomotor demands. It will also aid in the development of targeted treatments when musculotendon mechanics are altered by neuromuscular injuries or pathologies, and in the development and control of biomimetic assistive devices [8–10]. As such, we sought to determine how the relative contributions from the triceps surae and Achilles tendon to the stiffness of the ankle varied between posture and movement to determine what is driving the decrease in ankle stiffness during conditions more relevant to locomotion.

There are distinct differences in the mechanisms that generate muscle and tendon stiffness, which impact their mechanical response to movement. Tendons are passive, exhibiting non-linear, strain-dependent changes in stiffness. Although *in vivo* studies have demonstrated that the Achilles tendon exhibits complex viscoelastic behavior [11–13], it is commonly approximated as a non-linear spring with stiffness increasing throughout the toe region (up to about 30% of the force during a maximum voluntary contraction; MVC), after which it plateaus to a near-constant value [10, 14, 15]. Consistent with this assumption, Achilles tendon stiffness *in vivo* is often quantified within the linear region and reported as a constant value [11, 16–19]. Even when tendon stiffness has been quantified during human hopping, tendon stiffness was still represented as a single, constant value despite hysteresis being observed [13]. To our knowledge, there has been very little investigation into how tendon stiffness varies during more dynamic conditions aside from its well-documented hysteresis [11–13]. As such, it is unclear to what extent tendon stiffness varies between posture and movement.

Changes in both intrinsic and reflexive muscle stiffness between posture and movement could alter the contribution of muscle stiffness to the overall stiffness of the joint. Activation-dependent changes in muscle stiffness are thought to arise from the intrinsic stiffness of attached cross-bridges [20, 21] and from stretch-evoked reflexes that stiffen the muscle through a burst of muscle activation [22]. Movement can alter both the intrinsic and reflexive stiffness of muscle. Ettema and Huijing [23] demonstrated in rat muscle-tendon units that intrinsic muscle stiffness varies between isometric, concentric, and eccentric contractions, with stiffness being greatest during eccentric and lowest during concentric contractions. Additionally, reflexive stiffness can be altered volitionally, decrease during movement compared to posture, and can vary throughout the gait cycle [24, 25].

The purpose of this study was to investigate how the triceps surae and Achilles tendon contributions to ankle stiffness differ during conditions relevant to locomotion (i.e., movement) versus those relevant to stance (i.e., posture). We used our recently developed measurement technique that combines B-mode ultrasound imaging with joint-level perturbations to quantify ankle, muscle, and tendon stiffness simultaneously [26]. Plantarflexion torque was matched during the posture and movement conditions since ankle, muscle, and tendon stiffness are proportional to torque (or musculotendon force) [4, 14, 27, 28]. We also matched ankle angle, which is another factor contributing to ankle stiffness [29, 30]. Our central hypothesis was that the decrease in ankle stiffness during movement compared with posture is driven by a decrease in muscle stiffness. Secondary to our main objective, we sought to investigate the mechanisms leading to reduced joint stiffness during movement compared to posture. Thus, we evaluated if differences in muscle or tendon stiffness could be attributed to small differences in plantarflexion torque, short-latency stretch reflex activity, or muscle activation. These factors were evaluated because they can directly influence muscle and tendon stiffness. Collectively, our results identify the respective roles of the muscle and tendon in regulating ankle mechanics during conditions relevant to locomotion. Part of the data presented here has previously been used for a different purpose of tuning musculoskeletal models [10].

## METHODS

### Participants

Nine healthy young adults (age = 30 ± 3 years (mean ± standard deviation); height = 1.7 ± 0.1 m; body mass = 74 ± 16 kg; 5 males and 4 females) participated in this study. All participants were right-foot dominant and had no history of neuromuscular or musculoskeletal injuries to their right leg. Participants provided informed consent before participating. The Northwestern University Institutional Review Board approved all protocols (STU00009204 and STU00213839).

### Experimental setup

Participants were seated in an adjustable chair (Biodex Medical Systems, Inc. Shirley, NY). Their right leg was extended in front of them with a knee angle of 15° of flexion (Fig 1). A brace (Innovator DLX, Ossur, Reykjavik, Iceland) stabilized the knee in this position. The participant’s trunk and torso were stabilized with safety straps. We rigidly attached the participant’s right foot to an electric rotary motor (BSM90N-3150AF, Baldor, Fort Smith, AR) via a custom-made fiberglass cast. The cast extended distally from the medial and lateral malleoli to beyond the toes, creating a rigid foot that preserved the full range-of-motion of the ankle. We aligned the sagittal plane axis of rotation of the ankle to the center of rotation of the motor, restricting all movement to the sagittal plane. Throughout the experiment, we recorded ankle angle using a digital encoder integrated with the motor (24-bit, PCI-QUAD04, Measurement Computing, Norton, MA). A 6-degree-of-freedom load cell (45E15A4, JR3, Woodland, CA) measured all ankle forces and torques. We used xPC target (MATLAB, Mathworks, Natick, MA) to control the motor in real time. A position control scheme was used, so the motor dictated the position of the participant’s ankle at all times.

**Fig 1.**
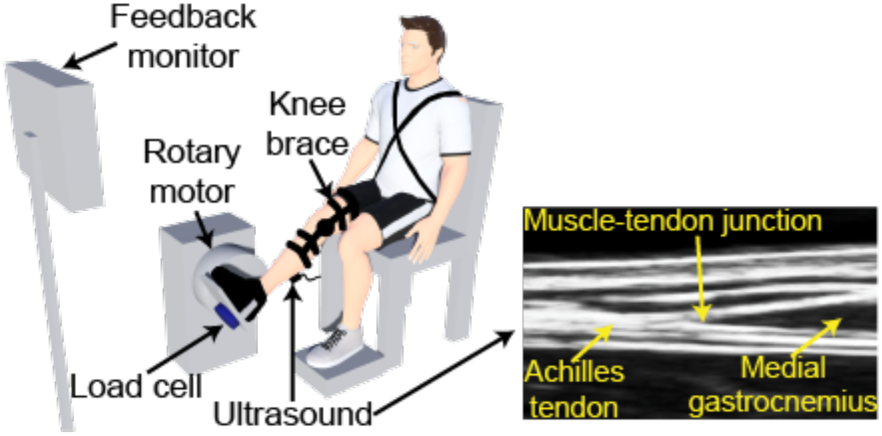
Schematic of the experimental setup. Participants sat with their foot secured to the rotary motor via a custom cast. The motor controlled the ankle angle, while the load cell measured the resultant torque; we simultaneously collected B-mode ultrasound over the muscle-tendon junction of the medial gastrocnemius. Figure adapted from [4].

We measured muscle activity from the medial gastrocnemius, lateral gastrocnemius, and soleus (ankle plantarflexors), and the tibialis anterior (ankle dorsiflexor) using single differential bipolar surface electrodes (Bagnoli, Delsys Inc, Boston, MA). We prepared the skin prior to electrode placement using standard procedures [31]. Electrodes were placed according to previously established guidelines [32]. Kinematic, kinetic, and EMG data were anti-aliased filtered at 500 Hz using a 5-pole Bessel filter and sampled at 2.5 kHz with a 24-bit data acquisition system (PCI-6289, Measurement Computing, Norton, MA, USA).

We used a B-mode ultrasound system to record images of the medial gastrocnemius muscle-tendon junction (MTJ) (Fig 1; Linear transducer LV7.5/60/128Z-2, LS128, CExt, Telemed, Lithuania). We have demonstrated previously that the results during active isometric contractions do not vary when imaging the various triceps surae muscles (medial gastrocnemius vs. lateral gastrocnemius vs. soleus) [26]. Therefore, we imaged the medial gastrocnemius muscle, which provided the highest quality image among the three muscles. The probe was positioned such that the MTJ was centered on the image. We secured the probe to the participant’s leg using a custom-made probe holder and elastic adhesive wrap (Coban™, 3M, St. Paul, MN). A trigger synchronized ultrasound data collection with all other measurements. Ultrasound images were acquired with a mean frame rate of 104 Hz and saved for offline processing.

### Protocol

At the start of the experiment, participants completed maximum voluntary contractions (MVCs). These data were used to normalize the EMGs [33] and to scale the visual feedback provided to the participants in later trials. Participants completed three MVC trials in both plantarflexion and dorsiflexion directions with the ankle angle fixed at 100°, each lasting 10 seconds.

Our primary objective was to determine how muscle, tendon, and ankle stiffness varied between posture and movement. Therefore, participants completed two conditions: 1) a movement condition, and 2) a posture condition (Fig 2). During the movement condition, the motor moved the participant’s ankle through a sinusoidal motion with an amplitude of 20° and a frequency of 0.5 Hz, resulting in periods of concentric and eccentric muscle contractions. The movement was centered at 100°. The sinusoidal ankle motion was chosen since it is similar to ankle kinematics during walking [34, 35]. Twenty-one movement trials were collected, each lasting 40 seconds.

**Figure 2.**
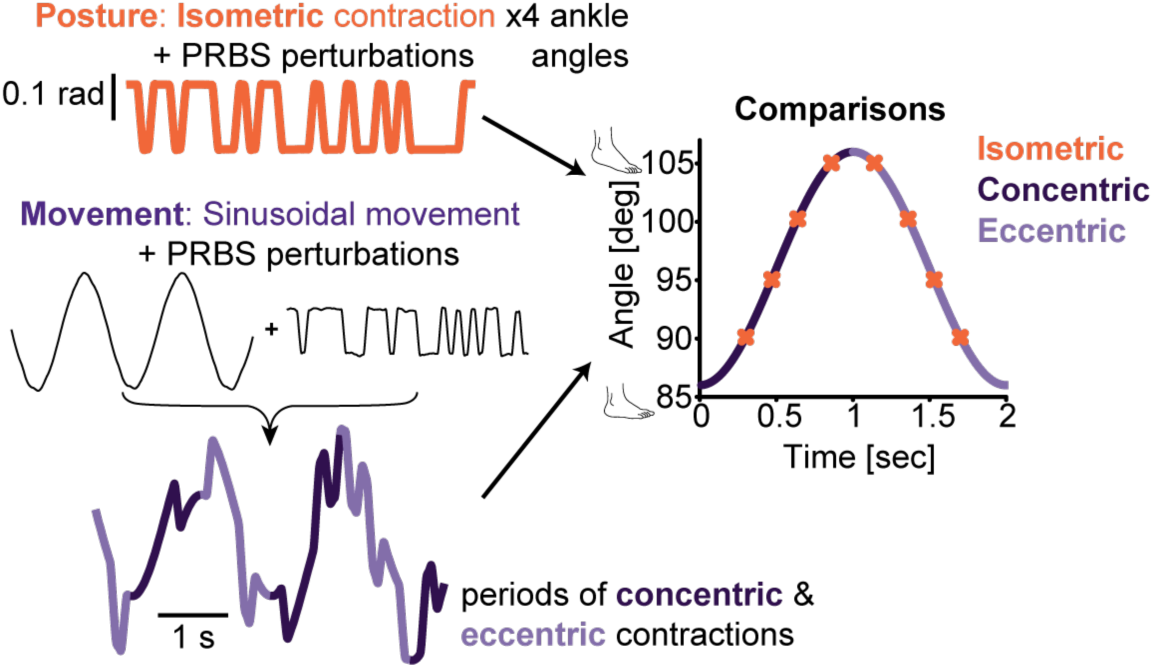
Experimental paradigm. To examine how muscle, tendon, and ankle stiffness differed between posture (orange) and movement (purple), participants completed two conditions: (1) a movement condition, and (2) a posture condition. During the movement condition (*bottom left*), the motor moved the participant’s ankle through a 20° sinusoidal motion at 0.5 Hz, resulting in periods of concentric (dark purple) and eccentric (light purple) contractions. PRBS perturbations were superimposed on the sinusoidal motion. Participants completed isometric contractions for the posture condition (*top left*) with the ankle angle at 90, 95, 100, and 105°. Therefore, all comparisons between concentric, eccentric, and isometric contractions were made at matched ankle angles (*right*).

This large number was needed for the time-varying system identification described below. For the posture condition, participants produced isometric contractions at four angles spanning much of the range of motion covered during the movement condition (90, 95, 100, and 105°). Three 40-second posture trials were collected at each ankle angle for a total of 12 trials.

Since ankle, muscle, and tendon stiffness are sensitive to torque or musculotendon force [4, 14, 27, 28], participants were instructed to sustain a constant plantarflexion torque of 15% MVC during each condition. We selected the magnitude of the target torque to be feasible to sustain without fatigue throughout the experiment. Real-time visual feedback of torque was provided to assist in this task. Tibialis anterior EMG feedback was also provided to help reduce co-contraction. For all conditions, practice was allowed so participants could become proficient with the task. Rest breaks were provided between each trial to prevent fatigue. Participants completed the conditions in a randomized fashion.

To estimate ankle, muscle, and tendon stiffness simultaneously, we used our experimental approach that combines controlled displacements of ankle angle, measures of ankle torque, and displacement of the medial gastrocnemius muscle-tendon junction (MTJ) from B-mode ultrasound. Thus, the rotary motor applied continuous random rotational perturbations during the postural conditions, and these perturbations were superimposed on the sinusoidal motion during the movement condition (Fig 2). The perturbations were pseudo-random binary sequence (PRBS) perturbations with an amplitude of 0.14 radians, a maximum velocity of 1.75 radians per second, and a switching time of 153 ms.

### Data processing and analysis

All data processing and signal analyses were performed using custom-written software in MATLAB. The same experimenter manually digitized the MTJ within each frame of the ultrasound video. Ultrasound data were synchronized with all other data [36] and linearly interpolated to the sampling rate of all other data (2.5 kHz).

### Ankle, muscle, and tendon stiffness

We used non-parametric system identification to compute ankle, muscle, and tendon impedance. Impedance is the dynamic relationship between an imposed displacement and the resultant force or torque [3]. The stiffness, or position-dependent component of impedance, is our primary metric of interest due to its relevance in the control of posture and movement [3, 37]. Ankle, muscle, and tendon stiffness were estimated from the measured ankle angle, ankle torque, and MTJ displacement (Fig 3). One challenge when estimating impedance during the movement condition is the low-frequency torque fluctuations that often occur when attempting to maintain a constant torque during the imposed sinusoidal movement. This low-frequency torque was removed by a second-order high-pass Butterworth filter with a cutoff frequency of 0.5 Hz. This cutoff frequency is below the range we previously showed is appropriate for estimating stiffness, our primary outcome measure [26]. While this was not necessary during the posture condition, we high-pass filtered the ankle angle, ankle torque, and MTJ displacement data for both conditions for consistency in processing. After this initial filtering, all data (ankle angle, ankle torque, and MTJ displacement) were decimated to 100 Hz.

**Fig 3.**
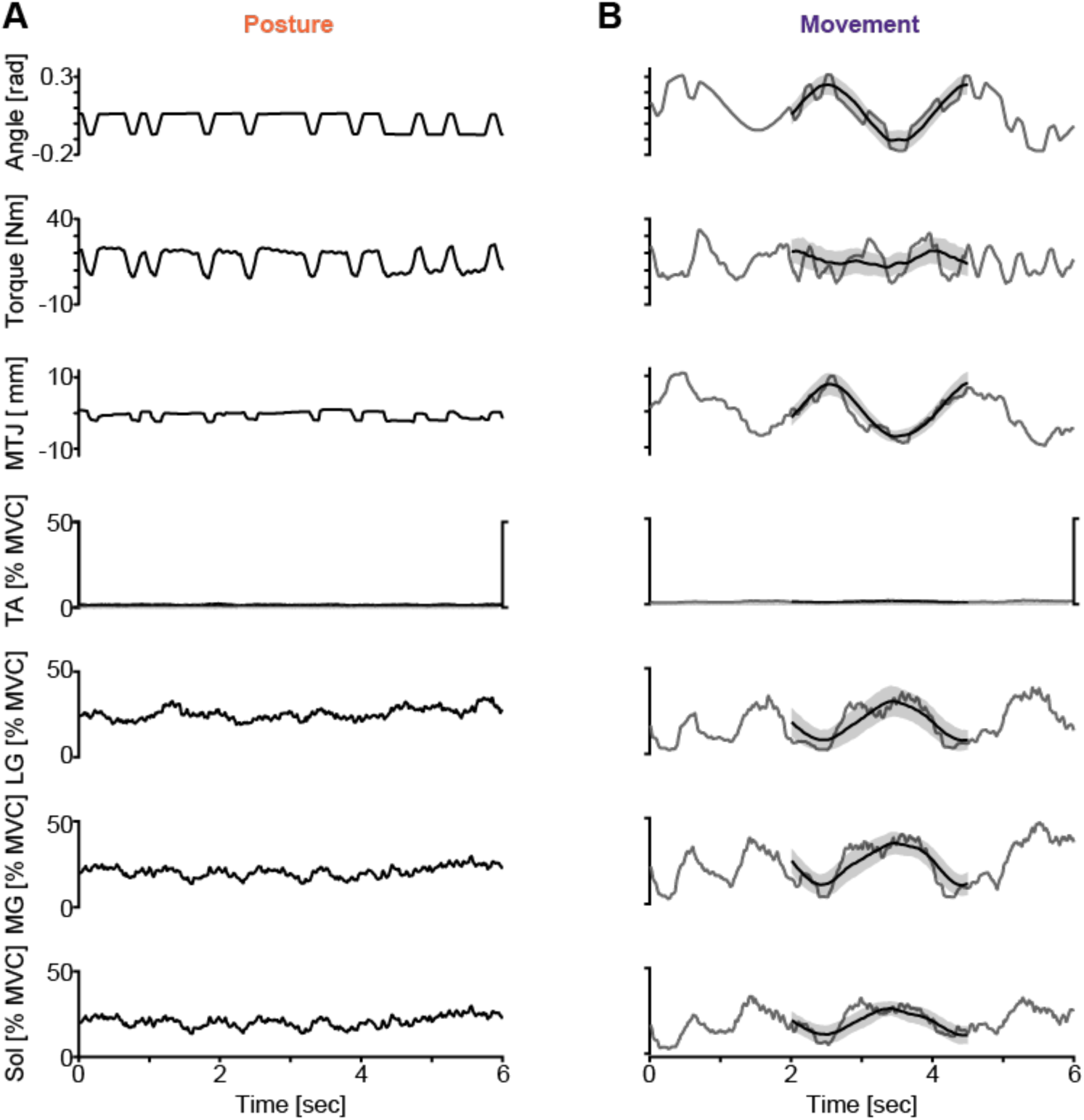
Sample data from a representative subject used to estimate ankle, muscle, and tendon stiffness during posture and movement conditions. For the posture condition (A), there are 6-second snippets of the imposed ankle angle, and measured ankle torque and muscle-tendon junction displacement. For the movement condition (B), there is a 6-second snippet from a single realization (gray) and the mean from the 200 realizations (black). The shaded black region is the standard deviation across the 200 realizations. These data were used to estimate ankle, muscle, and tendon stiffness. For both the posture and movement conditions, electromyography (EMG) data from the tibialis anterior (TA), lateral gastrocnemius (LG), medial gastrocnemius (MG), and soleus (Sol) are shown.

To calculate impedance during time-varying conditions (i.e., the movement condition), the system identification algorithm requires multiple repetitions of repeated data [38]. Therefore, all data from the movement condition were segmented into overlapping three-period long segments. Each segment started one period after the previous one. The realization was removed if the tibialis anterior was active (see *EMG Analysis* below). The tibialis anterior was deemed active if the activation within the realization exceeded 5% MVC. We used the 200 realizations where the plantarflexion torque produced was the most constant.

To quantify ankle, muscle, and tendon stiffness, we used our recently developed method [26]. Ankle impedance was quantified from the relationship between the imposed ankle rotations and the resultant ankle torque [3]. We assumed that the muscle and tendon are connected in series, and that muscle-tendon unit displacement can be determined by the rotation of the ankle multiplied by the Achilles tendon moment arm. We refer to the relationship between MTJ displacement and the angular rotations of the ankle as the translation ratio. Specifically, we used a non-parametric time-varying system identification algorithm [38]. While this was not necessary during the time-invariant posture condition, we used the same algorithm for both conditions to ensure consistent processing. The algorithm computed the time-varying impulse response functions (IRFs) for ankle impedance and the translation ratio at each time point along the movement profile during the movement condition, or at each ankle angle during the posture condition. The stiffness component of ankle impedance and the translation ratio for both the posture and movement conditions were computed by integrating the IRFs. From the estimates of ankle stiffness and the static translation ratio, we computed muscle and tendon stiffness algebraically [26].

We used a bootstrapping procedure to calculate the confidence intervals for our ankle, muscle, and tendon stiffness estimates during the movement condition. The 200 realizations were sampled randomly with replacement to produce a new ensemble of realizations. The new ensemble was then used to compute stiffness. We repeated this procedure 100 times, resulting in a distribution of stiffnesses. In our primary analysis, ankle, muscle, and tendon stiffness were normalized by the maximum plantarflexion torque to standardize our data across participants. As such, we refer to all stiffness estimates as the normalized stiffness. Ankle torque was also normalized by its maximum for comparisons across participants.

A single approximation of the Achilles tendon moment arm of 51.3 mm was used for all analyses. The moment arm was estimated as the mean across subjects from Clarke et al. [39] with an ankle angle of 100°. While the Achilles tendon moment arm does vary with ankle angle, the change in the moment arm across the tested angles is small, with a maximum difference of 4% compared with the moment arm at 100° [39]. We have previously evaluated the sensitivity of our muscle and tendon stiffness estimates to errors in the moment arm [26]. Over the tested torques, the sensitivity of muscle and tendon stiffness estimates to errors in the moment arm of ± 4% are relatively constant, and decreasing the error has little impact on the sensitivity. Lastly, it has been demonstrated that the Achilles tendon moment arm does not scale with anthropometric data [39, 40]. Therefore, we approximated the moment arm as a single value.

### EMG analysis

We examined the EMG data to determine: 1) if stiffness varied with mean muscle activity; and 2) if changes in reflex activity contributed to any observed difference in stiffness between posture and movement. All EMG data were notch-filtered to remove 60 Hz noise, demeaned, and rectified. We then processed EMG data in two ways. To determine if mean muscle activity varied between posture and movement and if that variation was related to differences in stiffness, the EMG signals were smoothed with a 25-ms moving average filter and decimated to 100 Hz. The EMG signals during the movement condition were segmented as described in *Ankle, muscle, and tendon stiffness*. All EMG signals were then normalized by the peak amplitude of the MVC trials (Fig 3). To examine the reflex activity, we quantified the perturbation-elicited responses in the triceps surae muscles (medial gastrocnemius, lateral gastrocnemius, and soleus). Only perturbations that lengthened the triceps surae were considered. For posture and movement trials, background activity was defined as the mean activity 40 ms prior to the perturbation, while short-latency reflex activity was defined as the mean activity from 35 ms to 60 ms following the onset of the perturbation [41]. To characterize the reflex response, background activity was subtracted from the short-latency stretch reflex activity. For the posture condition, data in each trial were segmented and aligned to the onset of each perturbation. The EMG segments were averaged, resulting in an average response from which the short-latency stretch reflex and background activity could be determined. For the movement condition, the perturbation-elicited responses within a 100 ms window at each time point along the movement profile were averaged, similar to the posture condition. This resulted in an estimate of the short-latency stretch reflex activity and background activity at each time point along the movement profile. We are interested in the reflex activity during the movement condition corresponding to the same joint angles tested during the posture condition. Thus, we compensated for sensorimotor processing delays by shifting the time windows of the background activity forward by 20 ms, and reflex activity shifted backward in time by 50 ms, so they are centered on the perturbation linking those quantities.

### Statistical Analysis

The goal of this study was to investigate how the triceps surae and Achilles tendon contributions to ankle stiffness differ between posture and movement. Participants completed isometric contractions in the posture condition at ankle angles of 90, 95, 100, and 105°. Therefore, all comparisons between posture and movement were made at matched ankle angles. Our central hypothesis was that a decrease in muscle stiffness primarily drives the decrease in ankle stiffness during movement. The movement condition resulted in periods of concentric and eccentric contractions (Fig 2), and differences in muscle stiffness between concentric and eccentric contractions have been observed previously [23]. We therefore separated the movement condition into concentric and eccentric contractions to determine whether contraction type also influenced muscle, tendon, or ankle stiffness. We used a linear mixed-effects model with stiffness as the dependent variable, ankle angle, contraction type (concentric, eccentric, and isometric), and structure (ankle, muscle, or tendon) were treated as fixed factors, while subject was a random factor. Interaction terms between angle, contraction type, and structure were also included. We performed all statistical analyses in MATLAB. For all models, we used a restricted maximum likelihood method to approximate the likelihood of the model, and Satterthwaite corrections for degrees of freedom [42]. Significance was set *a priori* at α=0.05. We used Holm post-hoc corrections for multiple comparisons. All reported p-values are the Holm-adjusted p-values [43]. All metrics reported are mean ± standard error unless otherwise noted.

## RESULTS

This study sought to determine how the relative contributions from the triceps surae and Achilles tendon to the stiffness of the ankle varied between posture and movement to determine what is driving the decrease in ankle stiffness during movement. First, we evaluated if there are differences in ankle, muscle, and tendon stiffness between posture and movement conditions. Second, we assessed how muscle and tendon stiffness contributions to ankle stiffness differ between posture and movement. Lastly, we investigated potential mechanisms contributing to changes in muscle and tendon stiffness between posture and movement, including differences in plantarflexion torque, muscle activation, and short-latency stretch reflex activity. While we made measurements at multiple ankle angles, our main findings were largely consistent across joint angles. Therefore, in the main text, we only describe the main effect of contraction type (e.g., comparing the isometric contractions (posture condition), concentric, and eccentric contractions (movement condition). Results on the interaction between contraction type and ankle angle are documented in the supplemental material.

### Ankle, muscle, and tendon stiffness decreased during movement compared to posture

Normalized ankle stiffness was significantly lower during movement compared to posture (Fig 4A). Specifically, normalized ankle stiffness decreased by 44% between isometric and concentric contractions (p < 0.001), and 34% between isometric and eccentric contractions (p = 0.008). Additionally, normalized ankle stiffness was 16% lower during concentric contractions compared to eccentric contractions, but this difference was not significant (p = 0.20). The difference in ankle stiffness between posture and movement is in accordance with previous findings [5–7].

**Fig 4.**
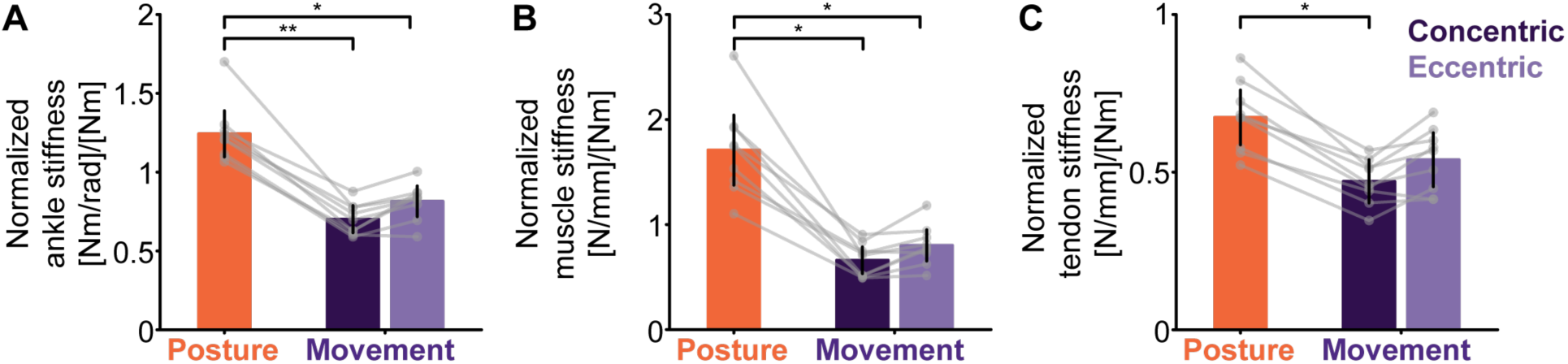
Normalized ankle, muscle, and tendon stiffness were significantly decreased during movement compared with posture. Mean normalized (*A*) ankle, (*B*) muscle, and (*C*) tendon stiffness for posture (isometric-orange) and movement (purple). The movement condition has been separated into concentric (dark purple) and eccentric contractions (light purple). Data are presented as the group mean ± 95% confidence interval from the linear mixed-effects model. The grey dots are individual subjects. Holm corrections were used for multiple comparisons; * indicates p<0.05, while ** indicates p<0.001.

Normalized muscle stiffness also decreased significantly during movement compared to posture (Fig 4B). Normalized muscle stiffness was significantly lower during both concentric and eccentric contractions compared with isometric contractions (concentric: 61% decrease; p = 0.004; eccentric: 53% decrease; p = 0.009). While not statistically significant, normalized muscle stiffness was 21% lower during concentric contractions compared to eccentric contractions (p = 0.29).

Normalized tendon stiffness also differed between posture and movement (Fig 4C), though to a lesser degree than muscle stiffness. Normalized tendon stiffness was 30% lower during concentric contractions compared to isometric contractions (p = 0.004). While not statistically significant, we observed a 20% decrease in normalized tendon stiffness during eccentric contractions compared to isometric contractions (p = 0.16), and a 15% decrease during concentric contractions compared to eccentric contractions (p = 0.84).

### Contributions from muscle and tendon to ankle stiffness differed between posture and movement

Next, we evaluated how the muscle and tendon contribute to ankle stiffness to determine what drives the decrease in ankle stiffness during movement. To do this, we evaluated if there was a significant difference between muscle and tendon stiffness within each condition (isometric, concentric, and eccentric). The muscle was 87% stiffer than the tendon during posture (p = 0.005; Fig 5A). In contrast, during concentric and eccentric contractions, while the muscle was 33% and 39% stiffer than the tendon, respectively, this did not reach statistical significance (p = 0.30 & p = 0.17, respectively).

**Fig 5.**
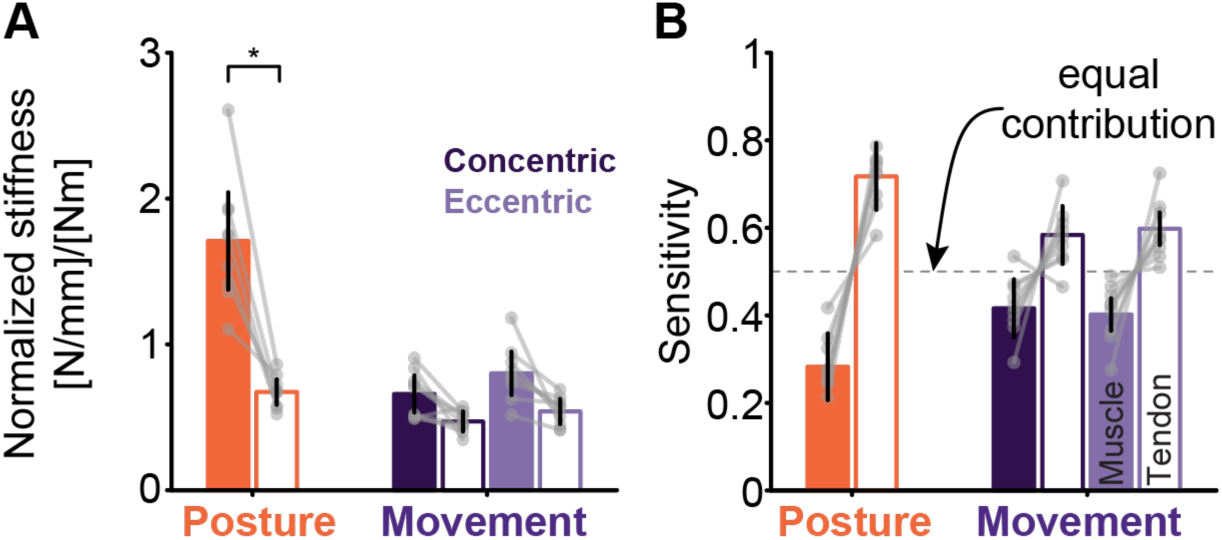
During posture, the muscle is significantly stiffer than the tendon, but during movement, muscle and tendon stiffness are more similar. This indicates a significant change in how the muscle (filled) and tendon (open) contribute to ankle stiffness. The movement condition has been separated into concentric (dark purple) and eccentric contractions (light purple). Data are presented as the group mean ± 95% confidence interval from the linear mixed-effects model. The grey dots are individual subjects. Holm corrections were used for multiple comparisons; * indicates p<0.05, while ** indicates p<0.001.

As an alternative way to evaluate the contribution of muscle and tendon stiffness to ankle stiffness, we quantified the sensitivity of ankle stiffness changes to changes in muscle or tendon stiffness within each condition. Ankle stiffness was 2.5 times more sensitive to changes in tendon stiffness than changes in muscle stiffness during postural conditions (Fig 5B). With the muscle and tendon connected in series, this suggests that the Achilles tendon is the dominant contributor to ankle stiffness during posture. During both concentric and eccentric contractions, ankle stiffness was 1.4 and 1.5 times more sensitive to changes in tendon stiffness than changes in muscle stiffness, respectively (Fig 5B). Thus, while the Achilles tendon is the dominant contributor to ankle stiffness during posture, the triceps surae and Achilles tendon contribute equally during movement, shifting ankle stiffness regulation from predominantly tendon-driven to a combined muscle and tendon mechanism.

### Confounding factors that can influence muscle and tendon stiffness

We sought to determine whether the observed differences in stiffness between posture and movement were attributed to confounding factors known to influence stiffness. To do so, we investigated if differences in plantarflexion torque, short latency stretch reflexes, or muscle activation could account for the observed stiffness differences between posture and movement.

#### Plantarflexion torque

While participants were instructed to maintain a constant level of plantarflexion torque, not all could do so during the movement task. As such, there were small differences in plantarflexion torque across contraction types (Fig 6). Plantarflexion torque was significantly lower during concentric contractions compared to isometric contractions (concentric: 11.7 ± 0.6% MVC; isometric: 15.2 ± 0.3% MVC; p = 0.001). It was also lower during concentric contractions compared with eccentric contractions (eccentric: 16.5 ± 0.8% MVC; p < 0.001). No significant difference was observed between isometric and eccentric contractions (p = 0.26). Since ankle, muscle, and tendon stiffness are sensitive to plantarflexion torque (or musculotendon force) [4, 14, 27, 28], we evaluated if the observed differences in torque could contribute to our stiffness results. This was done by using a previously established relationship between plantarflexion torque (or musculotendon force) and ankle, muscle, and tendon stiffness to correct for the small differences in ankle torque across the tested conditions [4]. We predicted stiffness using these models at the measured level of plantarflexion torque (or force) and divided our estimates of ankle, muscle, and tendon stiffness by these predicted values. We refer to these as the corrected stiffness. The differences in stiffness between posture and movement did not change when examining the corrected stiffness (Fig 7 and Table 1). These results indicate that the measured differences in plantarflexion torque do not account for the decrease in ankle, muscle, and tendon stiffness between posture and movement.

**Fig 6.**
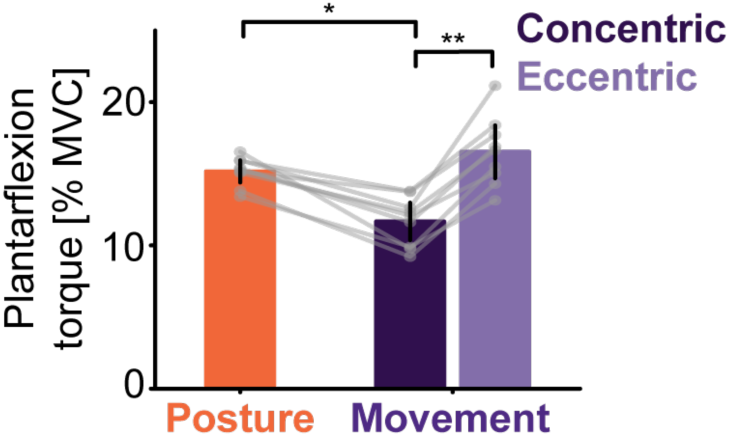
Plantarflexion torque significantly varied across contraction types. Mean ankle torque for posture (isometric-orange) and movement (purple). The movement condition has been separated into concentric (dark purple) and eccentric contractions (light purple). Data are presented as the group mean ± 95% confidence interval from the linear mixed-effects model. The grey dots are individual subjects. Holm corrections were used for multiple comparisons; * indicates p<0.05, while ** indicates p<0.001.

**Fig 7.**
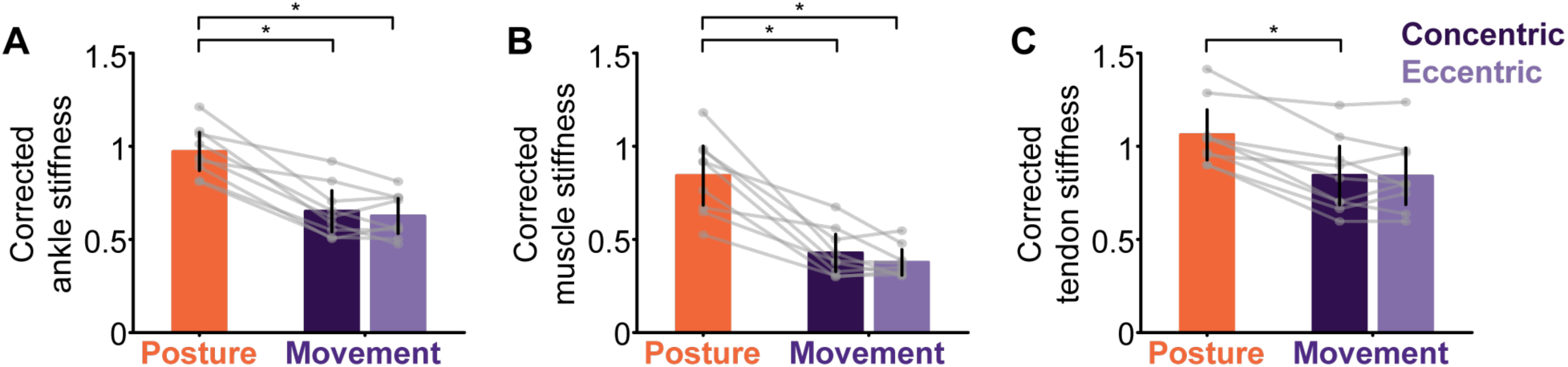
Corrected ankle, muscle, and tendon stiffness were significantly decreased during movement compared with posture. Mean corrected (*A*) ankle (*B*) muscle (*C*) and tendon stiffness for posture (isometric-orange) and movement (purple). Corrected stiffness values are our estimates of ankle, muscle, and tendon stiffness divided by model-predicted estimates. The movement condition has been separated into concentric (dark purple) and eccentric contractions (light purple). Data are presented as the group mean ± 95% confidence interval from the linear mixed-effects model. The grey dots are individual subjects. Holm corrections were used for multiple comparisons; * indicates p<0.05, while ** indicates p<0.001.

**Table 1.** Percent difference and Holm-adjusted p-values for all comparisons for corrected ankle, muscle, and tendon stiffness between isometric, concentric, and eccentric contractions.

|  | Isometric vs. Concentric | Isometric vs. Eccentric | Concentric vs Eccentric |
| --- | --- | --- | --- |
| Corrected ankle stiffness | 33%, $p = 0.007$ | 36%, $p = 0.004$ | 4%, $p = 1$ |
| Corrected muscle stiffness | 49%, $p = 0.02$ | 55%, $p = 0.009$ | 12%, $p = 1$ |
| Corrected tendon stiffness | 21%, $p = 0.02$ | 21%, $p = 0.07$ | 0.5%, $p = 1$ |

### Muscle activation

Our method assumes that changes in ankle torque are due to changes in the force transmitted through the Achilles tendon and that the dorsiflexor muscles are not active [26]. This was confirmed by measuring EMGs in the tibialis anterior (Fig 8), which was only 1 ± 0.03% MVC (mean ± standard deviation) across all tested conditions and all angles. There was also no significant difference in tibialis anterior activity across contraction types (Fig 8; all p > 0.94).

**Fig 8.**
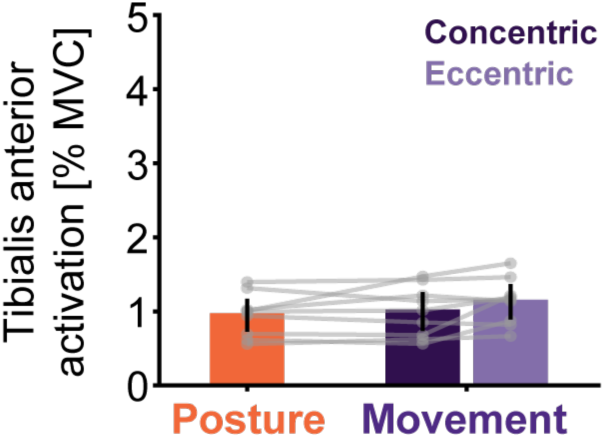
Muscle activity in the tibialis anterior did not vary between the posture and movement conditions. Tibialis anterior muscle activation for posture (isometric-orange) and movement (purple). The movement condition has been separated into concentric (dark purple) and eccentric contractions (light purple). Data are presented as the group mean ± 95% confidence interval from the linear mixed-effects model. The grey dots are individual subjects. Holm corrections were used for multiple comparisons; * indicates p<0.05, while ** indicates p<0.001.

We also examined if there were differences in muscle activation in the ankle plantarflexors between contraction types (Supplemental Fig 5). Differences in muscle activity were not observed between contraction types in the lateral gastrocnemius, medial gastrocnemius, or soleus. Thus, differences in plantarflexor muscle activation are unlikely to be the underlying mechanism causing the decrease in muscle stiffness during movement.

Stretch reflexes can alter muscle stiffness [22], and they can differ between posture and volitional movements [24, 25]. Even though our movements were imposed rather than volitional, we evaluated if there were differences in the stretch reflex across our tested conditions. We found that the short-latency stretch reflex did not vary between posture and movement (Fig 9 and Table 2). The short-latency stretch reflex in the medial gastrocnemius was 36% lower during eccentric contractions compared with concentric contractions (p = 0.02), but similar differences were not observed in the lateral gastrocnemius or soleus (Table 2), and all muscles contribute to our estimate of triceps surae stiffness [26]. Thus, it is unlikely that differences in stretch reflexes contributed to the observed decrease in muscle stiffness during movement.

**Fig 9.**
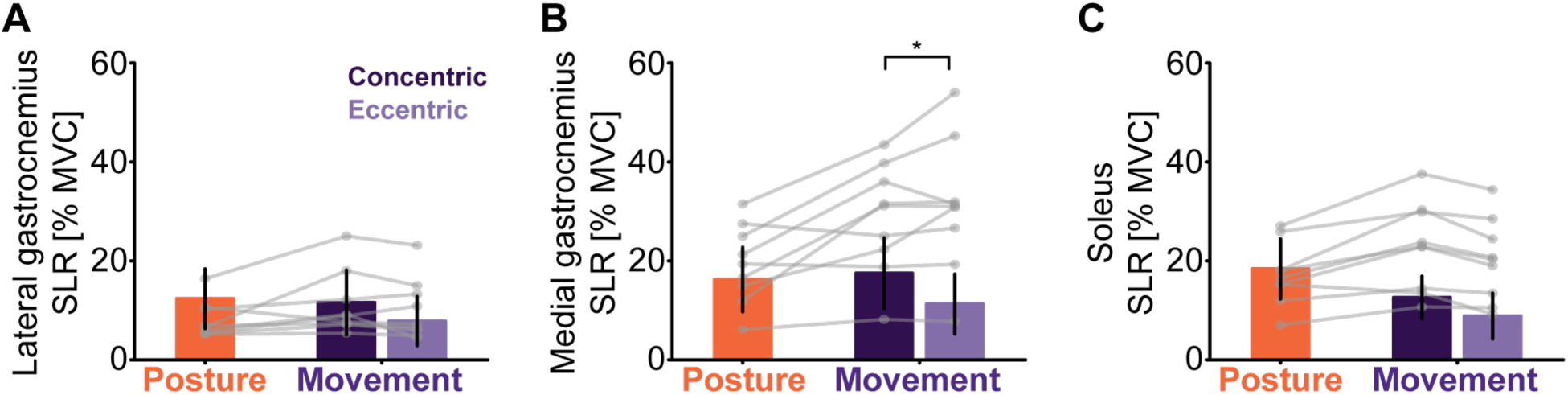
(A) lateral gastrocnemius (B) medial gastrocnemius (C) soleus short latency reflex (SLR) amplitude for posture (isometric-orange) and movement (purple). The movement condition has been separated into concentric (dark purple) and eccentric contractions (light purple). Data are presented as the group mean ± 95% confidence interval from the linear mixed-effects model. The grey dots are individual subjects. Holm corrections were used for multiple comparisons; * indicates p<0.05, while ** indicates p<0.001.

**Table 2.** Holm-adjusted p-values for all comparisons for the lateral gastrocnemius, medial gastrocnemius, and soleus short latency reflex response between isometric, concentric, and eccentric contractions.

|  | Isometric vs Concentric | Isometric vs Eccentric | Concentric vs Eccentric |
| --- | --- | --- | --- |
| Lateral gastrocnemius | $p = 1$ | $p = 0.81$ | $p = 0.53$ |
| Medial gastrocnemius | $p = 1$ | $p = 1$ | $p = 0.02$ |
| Soleus | $p = 1$ | $p = 0.27$ | $p = 1$ |

## DISCUSSION

This study quantified how triceps surae and Achilles tendon contributions to ankle stiffness differ between posture and movement. We observed that ankle, muscle, and tendon stiffness decreased during movement compared to torque- and angle-matched posture conditions. The decreases in muscle stiffness were substantially larger than the decreases in tendon stiffness, resulting in muscle and tendon stiffness being more similar during movement. Together, these findings reveal a shift in ankle stiffness from primarily dependent on tendon stiffness during posture to more equally dependent on muscle and tendon stiffness during movement.

### Mechanisms contributing to the decrease in muscle and tendon stiffness during movement

Importantly, the decrease in muscle stiffness during movement could not be attributed to differences in plantarflexion torque, muscle activation, or reflex activity; thus, we speculate that the reduction in stiffness reflects an intrinsic, history-dependent property of muscle, namely muscle thixotropy. During posture, our estimates of muscle stiffness are similar to measures of muscle short-range stiffness scaled to the triceps surae [26]. Muscle short-range stiffness describes the elastic muscle response to small, fast displacements [20, 21]. The amount of short-range stiffness within a muscle is also history-dependent, where movement can transiently reduce stiffness until it is held stationary again, also known as muscle thixotropy [44–48]. Much of our knowledge on history-dependent decreases in muscle stiffness comes from fiber-level paired-stretch paradigms where a conditioning stretch immediately precedes a test stretch [44, 49], or from sinusoidal oscillations [47] with limited investigation in whole muscle during continuous movements such as those tested here. It has been suggested that decreases in ankle stiffness induced by increasing postural sway were the result of muscle thixotropy [50]. However, given that Achilles tendon stiffness is the primary determinant of ankle stiffness during postural conditions [4, 51], and the lack of a direct measure of muscle and tendon stiffness, it is unclear what the source of the decrease in ankle stiffness with increased postural sway is. By directly quantifying triceps surae and Achilles tendon stiffness simultaneously during both posture and movement, our findings provide experimental evidence that movement is associated with a substantial reduction in muscle stiffness and supports the hypothesis that there are thixotropic decreases in muscle stiffness during movement.

Contrary to our hypothesis, tendon stiffness decreased during movement despite matched torque across conditions (Fig 4 & 7). Our hypothesis was rooted in the common assumption that the Achilles tendon is a non-linear spring such that stiffness scales with musculotendon force [10, 14, 15]. Nearly all *in vivo* estimates of Achilles tendon stiffness are made during fixed-end contractions, where participants are instructed to start from rest and contract maximally, resulting in no net change in muscle-tendon unit length, but changes in muscle and tendon length. Fixed-end contractions involve loading and unloading phases, but stiffness is commonly quantified as a single value, even when hysteresis is also quantified [11–13, 16–19]. To our knowledge, only one study has quantified Achilles tendon stiffness during both the loading and unloading phases of the fixed-end contraction and observed differences between the two phases, which is inevitable with hysteresis [52]. Moreover, no one has compared stiffness estimates under conditions where tendon length is held nearly constant, as tested here (Fig 3), and movement conditions. This matters because conditions where tendon length is nearly constant will limit hysteresis. Our findings provide experimental evidence that tendon mechanics are more complex than the commonly assumed non-linear spring and suggest that more complex representations that may include history dependence or other properties should be explored. While we observed movement-dependent changes in tendon stiffness, it is unclear what underlying mechanism drives the difference between posture and movement, and whether this relationship holds across different tasks.

### Limitations

There were a few limitations to our study. First, we tested our participants in a seated position using an imposed movement, which could impact the relevance of our estimates to volitional tasks like locomotion. While we used an imposed movement, a similar decrease in joint stiffness was observed during volitional movements [5, 6]. Additionally, our ankle stiffness values are similar to previously reported values of ankle stiffness during the early stance phase of walking [5, 53]. Our mean ankle stiffness values across all ankle angles were 57 ± 17 Nm/rad (mean ± standard deviation) or approximately 0.77 ± 0.23 (Nm/rad)/kg, during concentric contractions and 66 ± 21 Nm/rad (mean ± standard deviation) or approximately 0.89 ± 0.28 (Nm/rad)/kg during eccentric contractions. Lee and Hogan [53] report ankle stiffness values of approximately 56 ± 4 Nm/rad (mean ± standard error), while Rouse, et al. [5] reported ankle stiffness values of 1.5 Nm/rad/kg during early stance. Therefore, while we are in a more restricted setup, our values are similar to those in the literature.

Second, to exclude the possibility that small differences in plantarflexion torque contributed to our findings, we corrected ankle, muscle, and tendon stiffness by predicted values from an isometric model. The predicted values were based on previously established relationships between plantarflexion torque (or musculotendon force) and ankle, muscle, and tendon stiffness during isometric contractions in a similar experimental setup [4]. However, we note that there are a few differences between the two studies that could impact our results. First, the previous study was only performed with the ankle set at 90°; however, our current experiment tested ankle angles ranging from 90° to 105°. Ankle, muscle, and tendon stiffness during isometric contractions were similar across the tested ankle angles (Supplemental Fig 1). Therefore, differences in ankle angle likely had a minimal impact on our results. Second, the previously established relationship between torque (or musculotendon force) and stiffness was based on an isometric model, not on data during movement. The relationship between torque (or force) and stiffness may vary with contraction type, and this isn’t captured in this analysis. However, even if there are errors in the torque-stiffness relationship across contraction types, that resulting error would be relatively small compared to the observed decrease in muscle stiffness. Further, our main conclusions are drawn from the comparisons of muscle and tendon stiffness within each contraction type, which are unaffected by this secondary analysis.

While we use non-parametric system identification to compute ankle, muscle, and tendon impedance, we are reporting the stiffness component of those impedances. We have previously demonstrated, using non-parametric frequency-response functions (FRF), that stiffness is the dominant contributor to impedance at the frequencies at which we have power (approximately 1-6.5 Hz) [4, 26, 54]. While we are estimating impedance and stiffness via IRFs, during the posture condition, ankle stiffness and the static translation ratio estimated via IRF were similar to estimates from FRF (ankle: mean difference = 1.25 (Nm/rad)/Nm or 2% of the measured value; static translation ratio: 0.007 or 2% of the measured value). Moreover, both FRFs and IRFs described the data well (ankle VAF: FRF = 95%; IRF = 93%; static translation ratio VAF: FRF = 87% and IRF = 81%), and are consistent with our previously reported values [4, 26, 54]. Thus, stiffness is a good representation of the observed behavior of the ankle, muscle, and tendon.

### Applications and conclusions

We are the first to present *in vivo* measurements of the muscle and tendon contributions to joint stiffness during movement, but understanding the musculotendon contribution to joint mechanics is a growing area of interest. Others have used musculoskeletal models to determine how muscle and tendon stiffness contribute to the net mechanics of a joint [8, 55]. While the model-based approaches are a good first approximation, they can provide unrealistic estimates of muscle and tendon stiffness [10]. Thus, we have proposed a combined approach in which musculoskeletal models can be calibrated using experimental measures [10]. These models can simulate a broad range of conditions, including those where our system identification-based approach cannot be applied. Continued development and validation of experimentally informed musculoskeletal models will be critical for accurately capturing the dynamic and context-dependent contributions of muscle and tendon to joint mechanics across a wide variety of functionally relevant human movement, and in situations when movement is impaired due to aging, injury, or disease.

Our findings establish that decreases in ankle stiffness during movement compared with posture result largely from decreases in muscle stiffness. These large changes in muscle properties alter the respective roles of the muscle and tendon between posture and movement; the triceps surae and Achilles tendon contribute more equally to the stiffness during movement. These results provide fundamental insight into how ankle mechanics are governed, which is critical for understanding humans’ ability to navigate their physical world and may help guide rehabilitation when neuromuscular pathologies alter ankle mechanics.

## CONFLICT OF INTEREST

The authors declare no competing interests.

## AUTHOR CONTRIBUTIONS

KLJ, DL, SSML, and EJP conceived of the study and designed the experimental protocol and analyses. KLJ carried out the experiments, analyzed the data, and drafted the manuscript. KLJ, DL, SSML, and EJP edited the manuscript. All authors approved the final version.

## FUNDING

Research reported in this publication was supported by the National Institute On Aging of the National Institutes of Health under Award Number F31 AG069412. The content is solely the responsibility of the authors and does not necessarily represent the official views of the National Institutes of Health. Research supported by the American Society of Biomechanics’ Graduate Student Grant-in-aid.

## DATA AVAILABILITY

The data from the current study are available from the corresponding author upon reasonable request.

## SUPPLEMENTAL RESULTS

### Ankle, muscle, and tendon stiffness variation with contraction type and ankle angle

**Supplemental Fig 1.**
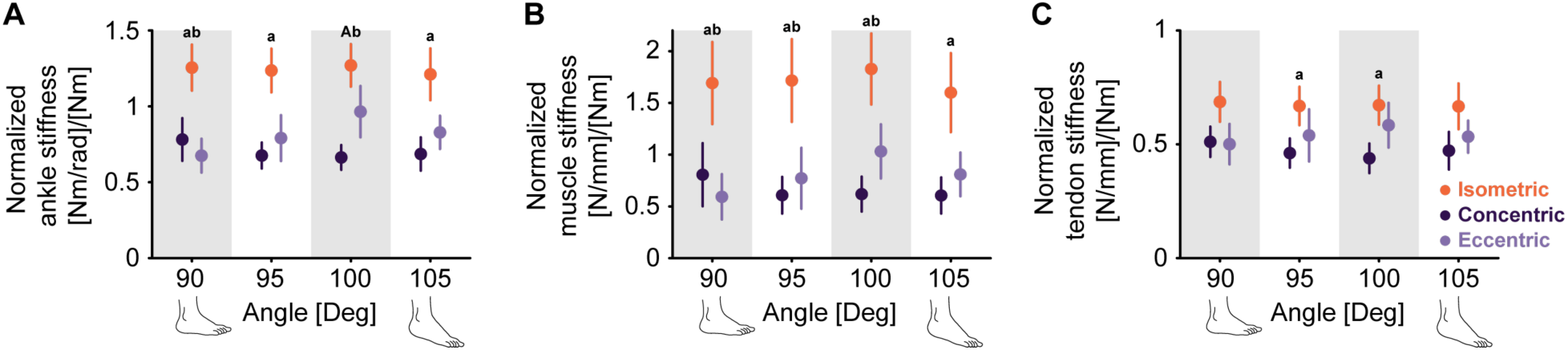
Mean normalized (*A*) ankle (*B*) muscle (*C*) and tendon stiffness for isometric (posture-orange), concentric (dark purple), and eccentric (light purple) contractions at each ankle angle. Data are presented as the group mean ± 95% confidence interval from the linear mixed-effects model. The letters indicate if stiffness varied significantly between contraction types: (a) isometric and concentric, (b) isometric and eccentric, and (c) concentric and eccentric contractions. Holm corrections were used for multiple comparisons; a capital letter indicates p<0.001, while a lowercase letter indicates p<0.05.

**Supplemental Fig 2.**
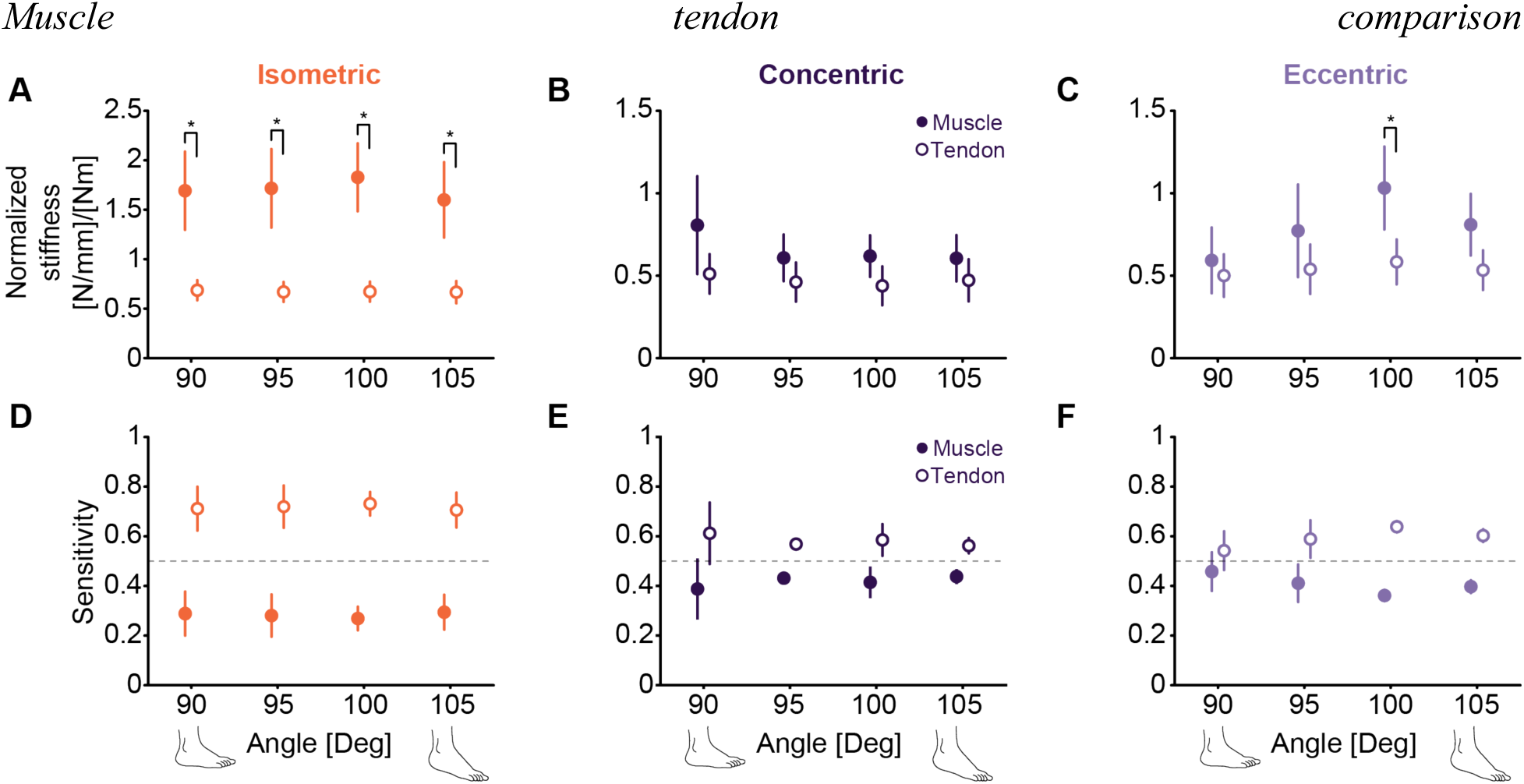
Comparison of muscle (filled) and tendon (open) during (*A*) isometric, (*B*) concentric, and (*C*) eccentric contractions. Sensitivity of ankle stiffness to changes in muscle stiffness and tendon stiffness during (*D*) isometric, (*E*) concentric, and (*F*) eccentric contractions. Data are presented as the group mean ± 95% confidence interval from the linear mixed-effects model. Holm corrections were used for multiple comparisons; * indicates p<0.05, while ** indicates p<0.001.

### Plantarflexion torque variation with contraction type and ankle angle

**Supplemental Fig 3.**
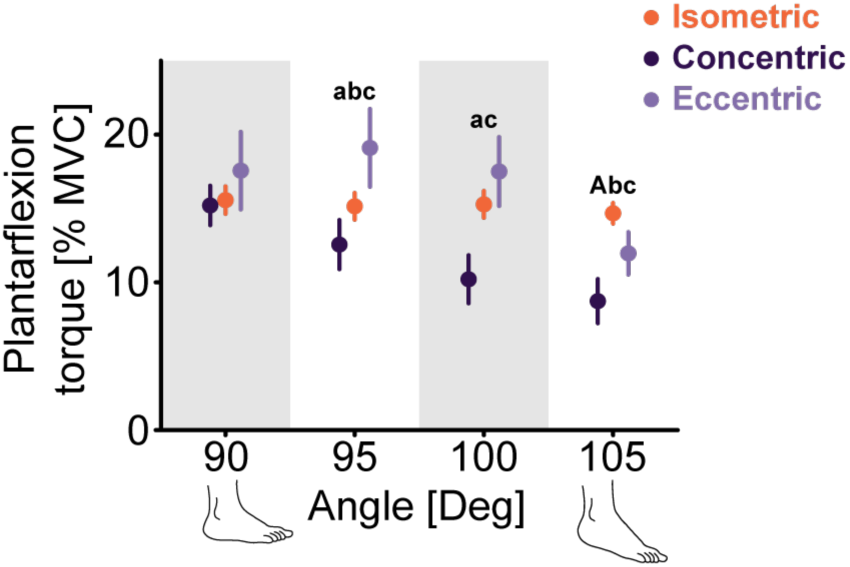
Mean plantarflexion torque for isometric (posture-orange), concentric (dark purple), and eccentric (light purple) contractions at each ankle angle. Data are presented as the group mean ± 95% confidence interval from the linear mixed-effects model. The letters indicate if stiffness varied significantly between contraction types: (a) isometric and concentric, (b) isometric and eccentric, and (c) concentric and eccentric contractions. Holm corrections were used for multiple comparisons; a capital letter indicates p<0.001, while a lower-case letter indicates p<0.05.

### Correct ankle, muscle, and tendon stiffness

**Supplemental Fig 4.**
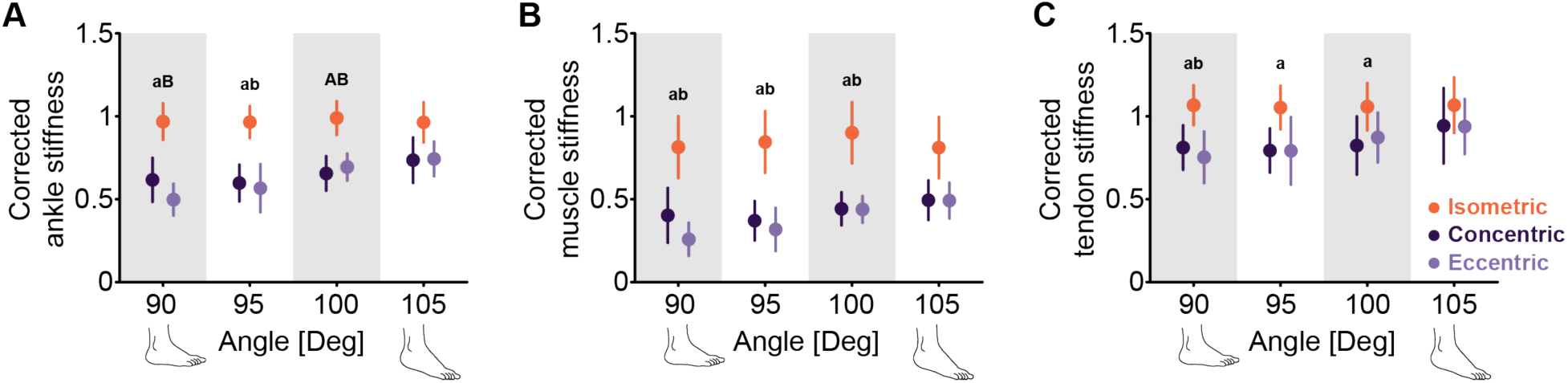
Corrected (*A*) ankle (*B*) muscle (*C*) and tendon stiffness for isometric (posture-orange), concentric (dark purple), and eccentric (light purple) contractions at each ankle angle. Data are presented as the group mean ± 95% confidence interval from the linear mixed-effects model. The letters indicate if stiffness varied significantly between contraction types: (a) isometric and concentric, (b) isometric and eccentric, and (c) concentric and eccentric contractions. Holm corrections were used for multiple comparisons; a capital letter indicates p<0.001, while a lowercase letter indicates p<0.05.

### Variation in plantarflexor muscle activation with contraction type

**Supplemental Fig 5.**
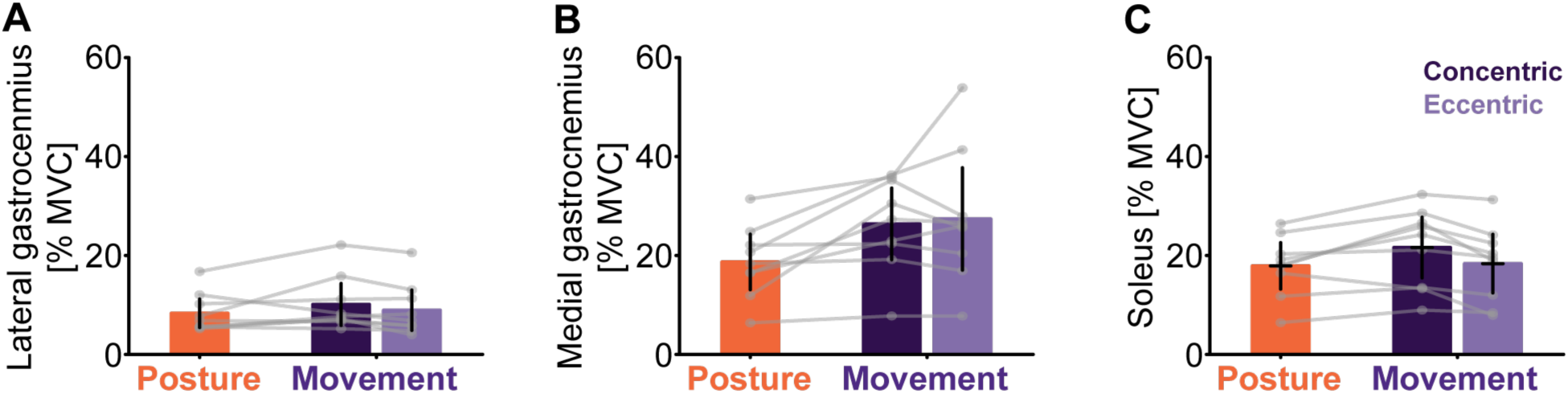
Muscle activity in the plantarflexors did not vary between posture and movement. (A) lateral gastrocnemius, (B) medial gastrocnemius, (C) soleus muscle activation for posture (isometric-orange) and movement (purple). The movement condition has been separated into concentric (dark purple) and eccentric contractions (light purple). Data are presented as the group mean ± 95% confidence interval from the respective linear mixed-effects models. Holm corrections were used for multiple comparisons; * indicates p<0.05, while ** indicates p<0.001.

**Supplemental Fig 6.**
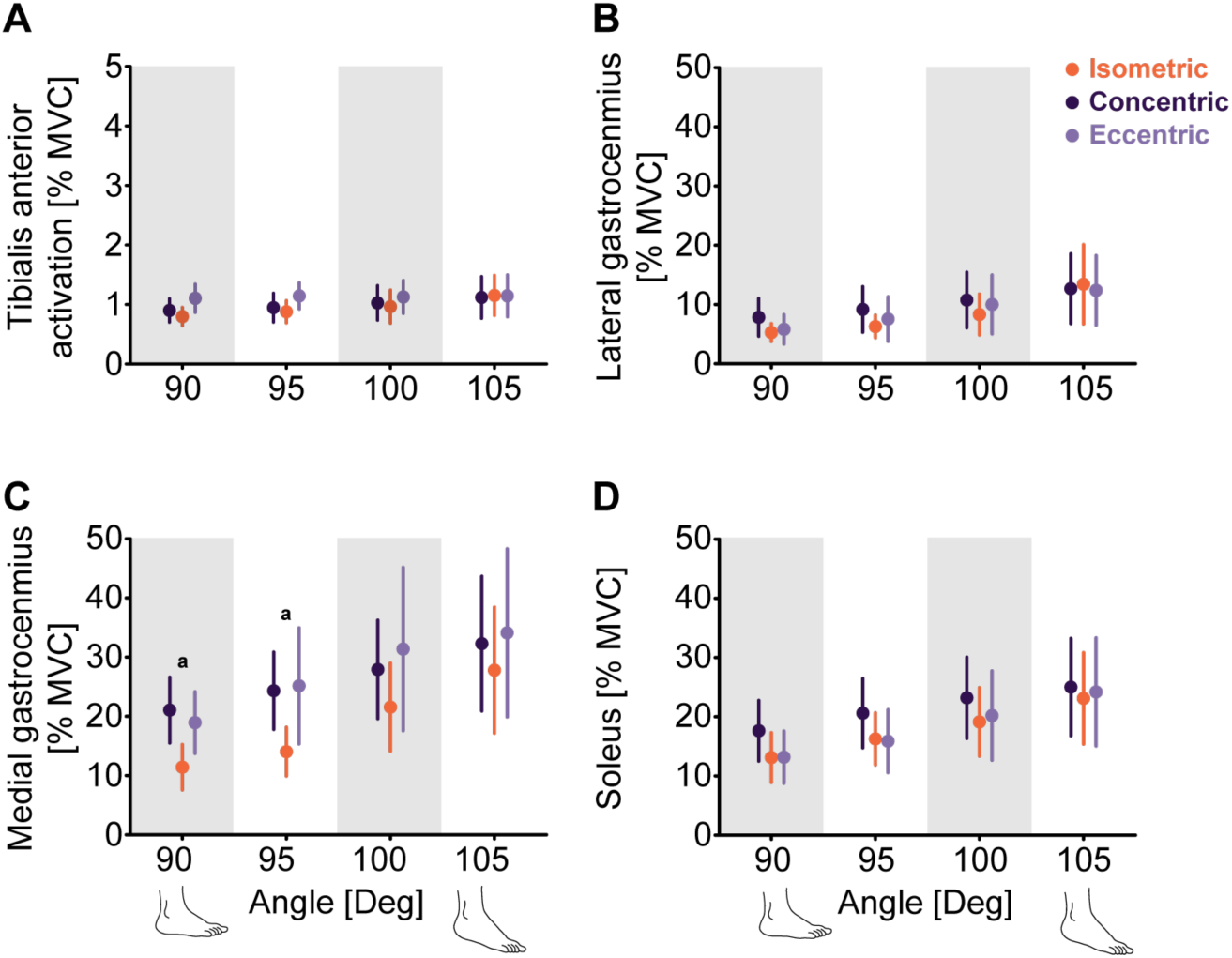
(A) Tibialis anterior (B) lateral gastrocnemius, (C) medial gastrocnemius, and (D) soleus muscle activation for posture (isometric-orange) and movement (purple). The movement condition has been separated into concentric (dark purple) and eccentric contractions (light purple). Data are presented as the group mean ± 95% confidence interval from the linear mixed-effects model. The letters indicate if stiffness varied significantly between contraction types: (a) isometric and concentric, (b) isometric and eccentric, and (c) concentric and eccentric contractions. Holm corrections were used for multiple comparisons; a capital letter indicates p<0.001, while a lowercase letter indicates p<0.05.

**Supplemental Fig 7.**
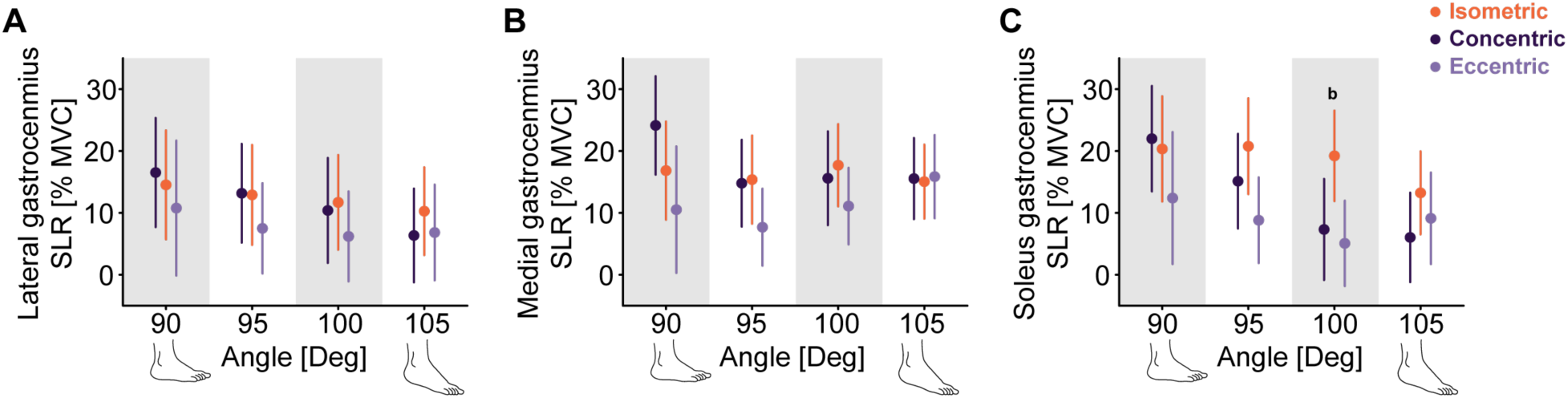
(A) Lateral gastrocnemius, (B) medial gastrocnemius, and (C) soleus muscle short-latency reflex (SLR) activation for posture (isometric-orange) and movement (purple Mean normalized (*A*) ankle (*B*) muscle (*C*) and tendon stiffness for isometric (posture-orange), concentric (dark purple), and eccentric (light purple) contractions at each ankle angle. Data are presented as the group mean ± 95% confidence interval from the linear mixed-effects model. The letters indicate if stiffness varied significantly between contraction types: (a) isometric and concentric, (b) isometric and eccentric, and (c) concentric and eccentric contractions. Holm corrections were used for multiple comparisons; a capital letter indicates p<0.001, while a lowercase letter indicates p<0.05.

## Notes

### Competing Interest Statement

The authors have declared no competing interest.

